# Preserved cognition in elderly with intact rhinal cortex

**DOI:** 10.1101/2022.05.30.494074

**Authors:** Farshid Sepehrband, Kirsten M. Lynch, Andrea Sotelo Gasperi, Michael S. Bienkowski, Xinhui Wang, Helena C. Chui, Arthur W Toga, the Alzheimer’s Disease Neuroimaging Initiative

**Affiliations:** Stevens Neuroimaging and Informatics Institute, Keck School of Medicine, University of Southern California, Los Angeles, California, USA; Alzheimer’s Disease Research Center, Keck School of Medicine, University of Southern California, Los Angeles, CA, USA; Dornsife College of Letters, Arts and Science, University of Southern California, Los Angeles, California, USA; Department of Neurology, Keck School of Medicine, University of Southern California, Los Angeles, CA, USA

## Abstract

Alzheimer’s disease pathology leads to neurodegeneration within the memory-related structures of the medial temporal cortex and hippocampus. Neurodegeneration also occurs as a part of normative aging and it is unclear whether medial temporal lobe subregions are selectively intact in older adults with preserved cognitive function in comparison to adults who are cognitively impaired. In this study, we used T1-weighted and high-resolution T2-weighted magnetic resonance images to assess age-related volumetric changes to medial temporal lobe regions, including the hippocampal formation and rhinal cortex, in patients with mild cognitive impairment and cognitively normal controls in two independent cohorts. Our results show age was significantly associated with regional atrophy in the hippocampus, but not the rhinal cortex. Additionally, variability in regional medial temporal lobe volume was associated with tau uptake in the rhinal cortex, but not the hippocampus. Together, these results suggest that the rhinal cortex may be more indicative of Alzheimer’s disease pathology and can help differentiate from age-related neurodegeneration.

## Introduction

Alzheimer’s disease (AD) is a progressive neurodegenerative disorder characterized by significant memory impairments that constitutes the most common form of dementia (Hebert et al., 2013). The hallmark pathological changes in AD, including deposition of tau neurofibrillary tangles and senile plaques, are associated with neuronal death and subsequent tissue loss in targeted brain structures (Arnold et al., 1991; Bejanin et al., 2017; Braak and Braak, 1991; Price et al., 2001). It is well known that these changes begin preferentially in the memory-related structures of the medial temporal lobe, including the hippocampus and rhinal cortex (Devanand et al., 2012; Pihlajamaki et al., 2009), and contribute to the pronounced cognitive deficits that define AD (Kerchner et al., 2012). Prior to the formal onset of AD, the transitional state between normal aging and dementia is known as mild cognitive impairment (MCI) (Cheng et al., 2017). In patients with MCI, MTL atrophy is a strong predictor for the conversion to AD (Apostolova et al., 2006; Devanand et al., 2007; Killiany et al., 2002; Stoub et al., 2005). Because patients with MCI have not yet developed the deleterious symptoms that clinically define AD, it is therefore important to characterize these initial structural changes for the development of early diagnostic biomarkers.

The hippocampal formation consists of several functionally specialized subfields with distinct cellular characteristics and includes the cornu ammonis (CA) 1-4, dentate gyrus (DG), and subiculum (SUB) (Duvernoy et al., 2013). The rhinal cortex envelopes the anterior part of the hippocampal formation and includes several cytoarchitecturally defined and interconnected cortical areas – the entorhinal cortex (EnRC), which receives multimodal sensory information from the perirhinal (PRC) and parahippocampal cortex (PHC), provides the major input to the hippocampal formation (Schultz et al., 2015). There is increasing evidence that MTL subregions are differentially vulnerable to pathological insults and follow hierarchical atrophic trajectories over the course of AD disease progression (Chételat, 2018; de Flores et al., 2015a). Previous neuropathological studies have demonstrated that tau first collects in the entorhinal cortex before spreading to the SUB and hippocampal CA regions (Arnold et al., 1991; Bobinski et al., 1995; Schönheit et al., 2004). Therefore, efforts to characterize clinically significant structural alterations in MCI and early AD benefit from considering MTL subregions separately.

Recent advances in neuroimaging have allowed for the characterization of individual MTL subregions in clinical populations (Iglesias et al., 2015; Winterburn et al., 2013; Yushkevich et al., 2015). The hippocampus has been the subject of many studies on early structural alterations in AD because hippocampal atrophy is one of the most marked macroscopic features in AD and has been reported in prodromal patients with MCI (de Flores et al., 2015a; Mak et al., 2017a; Ruan et al., 2016). Several studies have shown the earliest volumetric reductions occur predominantly in the SUB and CA1 subfields (Blanken et al., 2017; Gunten et al., 2006; Mak et al., 2017b; Padurariu et al., 2012; Su et al., 2018); however discrepant findings have also been reported with volume loss in CA2 (Jessen et al., 2010), presubiculum (Carlesimo et al., 2015), CA4 and DG subfields (Broadhouse et al., 2019) or sparing of hippocampal subfield volumes (Geoffrey A. Kerchner et al., 2013; Wisse et al., 2014). Unlike the widely studied hippocampal pattern of atrophy in AD, there has not been much research exploring atrophic patterns of other MTL subregions, even though previous research suggests that atrophy of the EnRC precedes that of the hippocampus (Du, 2001; Pennanen et al., 2004). Histological evidence in post-mortem samples show tau tends to collect first in the EnRC before spreading to the hippocampus and neocortex (Arnold et al., 1991; Bobinski et al., 1995; Schönheit et al., 2004), resulting in layer-specific decreases in neuron number (Gómez-Isla et al., 1996; Kordower et al., 2001).

Neurodegeneration also occurs as a part of normative aging. Brain shrinkage selectively affects the MTL regions (Raz and Rodrigue, 2006; Sowell et al., 2003) and is accompanied by age-related changes in cognition (Fjell & Walhovd 2010; Lockhart & DeCarli 2014). Because the difference between typical and disordered aging is not clear and the main risk factor for developing AD is advancing age (Fjell et al., 2014), it is challenging to confidently conclude whether an imaging-derived alteration reflects pathophysiology or is part of the normative neurodegenerative process during aging. There have been multiple studies looking at the MTL structures that are affected during normal aging, however results are inconclusive due to variability in the general population and differences in methods used (de Flores et al., 2015a). Several studies have found age-related volumetric reductions in CA1(Mueller et al., 2007; Raz et al., 2014; Shing et al., 2011), while others have found a selective diminution of SUB (de Flores et al., 2015c; La Joie et al., 2010; Ziegler et al., 2011), DG (Mueller and Weiner, 2009; Pereira et al., 2014; Wisse et al., 2014) or EnRC (G A Kerchner et al., 2013). It would therefore be valuable to know what MTL subregions are selectively intact in older adults with preserved cognitive function to compare these findings with the brain scans of older adults who are cognitively impaired.

In the present study, we sought to characterize the neurodegenerative trajectory of MTL sub-regions, including the hippocampus and rhinal cortex, in order to differentiate between age-dependent selective vulnerabilities to normative aging and early cognitive decline. We used data from the latest phase of the ADNI effort (ADNI-3 cohort) (Weiner et al., 2017) that consists of more than 600 cognitively normal controls and patients with amnestic MCI. T1-weighted and high resolution T2-weighted images were used to segment MTL sub-regions, including hippocampal subfields and rhinal cortical areas, and the age-related volumetric changes of these regions were quantified. In order to confirm our findings, we replicated the findings of this study in a second independent dataset of nearly 200 subjects from University of Southern California Alzheimer’s Disease Research Center (USC ADRC). Furthermore, we validated the pathological relevance of our findings with *in vivo* measures of regional neurofibrillary tangle deposition using PET imaging of tau uptake on the ADNI-3 cohort.

## Method

### Participants

Data used in the preparation of this article were obtained from the Alzheimer’s Disease Neuroimaging Initiative 3 (ADNI-3) database (http://adni.loni.usc.edu) (Weiner et al., 2017). ADNI was launched in 2003 as a public-private partnership, led by Principal Investigator Michael W. Weiner, MD. The primary goal of ADNI has been to test whether serial MRI, positron emission tomography (PET), other biological markers, and clinical and neuropsychological assessment can be combined to measure the progression of mild cognitive impairment (MCI) and early Alzheimer’s disease (AD). Results were then validated on data from University of Southern California Alzheimer’s disease research center (USC ADRC).

Statistical analysis was performed on a subset of the Alzheimer disease neuroimaging initiative 3 (ADNI-3) cohort (Weiner et al., 2017), in which high resolution T2-weighted turbo spin echo (T2w-TSE) hippocampal images were available. Data of 645 subjects with T1-weighted (T1w) and T2w-TSE MRIs were downloaded from the ADNI database (http://adni.loni.usc.edu) (Toga and Crawford, 2010). 41 participants were excluded after quality control, as described below in the *MRI analysis* section.

For replication data participants were recruited through the USC Alzheimer’s Disease Research Center (ADRC): combined USC and the Huntington Medical Research Institutes (HMRI), Pasadena, CA. We included 175 participants (**Table 2**) with T1w and high-resolution hippocampal images.

### Magnetic resonance imaging

MRI imaging of the ADNI-3 was done exclusively on 3T scanners (Siemens, Philips and GE) using a standardized protocol. 3D T1w with 1mm^3^ resolution was acquired using an MPRAGE sequence (on Siemens and Philips scanners) and FSPGR (on GE scanners). For T2w-TSE images, a 2D sequence with high in-plane resolution (0.4 × 0.4 mm^2^), with slice thickness of 2mm was acquired, which enabled medial temporal lobe (MTL) subfield segmentation. MPRAGE T1w MRI scans were acquired using the following parameters: Sagittal slices, TR = 2300 ms, TE = 2.98 ms, FOV = 240 × 256 mm^2^, matrix = 240 × 256 (variable slice number), TI = 900 ms, flip angle = 9, effective voxel resolution = 1 × 1 × 1 mm^3^. The FSPGR sequence was acquired using: Sagittal slices, TR = 7.3 ms, TE = 3.01 ms, FOV = 256 × 256 mm^2^, matrix = 256 × 256 (variable slice number), TI = 400 ms, flip angle = 11, effective voxel resolution = 1 × 1 × 1 mm^3^. The same standardized sequence was used across USC ADRC cohort. T2w-TSE sequence includes: Coronal 2D slices with 2mm thickness, flip angles = 122, matrix = 448 × 448 × 30, effective voxel resolution = 0.4 × 0.4 × 2 mm^3^, TR = 8020 ms, TE = 50 ms.

### MRI analysis

#### Preprocessing and parcellation

After downloading the raw images, *dcm2nii* was used to convert the *dicom* images to the *nifti* file format (Li et al., 2016). T1w preprocessing and parcellation was done using the freely available FreeSurfer (v5.3.0) software package (Fischl, 2012) and data processing was performed using the Laboratory of Neuro Imaging (LONI) pipeline system (http://pipeline.loni.usc.edu) (Dinov et al., 2010, 2009; Moon et al., 2015; Torri et al., 2012), similar to (Sepehrband et al., 2018a; Sta Cruz et al., 2019). Total brain, hippocampus, rhinal cortex (called entorhinal cortex in FreeSurfer atals), fusiform gyrus and parahippocampal cortex were segmented as part of the *recon-all* module of the FreeSurfer, which uses an atlas-based parcellation approach to segment subcortical regions (Desikan et al., 2006; Fischl et al., 2002). Prior to parcellation, *recon-all* applies the following pre-processing steps: motion correction, non-uniform intensity normalization, Talairach transform computation, intensity normalization and skull stripping (Dale et al., 1999; Desikan et al., 2006; Fischl et al., 2004b, 2004a, 2002, 1999; Fischl and Dale, 2000; Reuter et al., 2012, 2010; Reuter and Fischl, 2011; Segonne et al., 2007, 2004; Sled et al., 1998; Waters et al., 2018).

#### MTL subfield segmentation

MTL subfield volumes were generated by applying the automated segmentation of hippocampus subfield (ASHS) software to the hippocampal T2-weighted MRI scan (version 1.0.0), using T1w and T2w-TSE images (Yushkevich et al., 2015). ASHS uses a multi-atlas label fusion technique and a learning-based error correction module to segment hippocampal subfields, and MTL cortical regions. Segmentations are done along the entire length of the hippocampal formation. Segmentations are obtained using high-dimensional mapping to multiple manually labeled atlas images and then fused into a consensus segmentation, taking into account the degree of similarity between the subject image and atlas images [ref]. Patterns of systematic segmentation errors introduced in this procedure are learned *a priori* using training data for postprocessing correction to generate final segmentation. ASHS segments the following regions: dentate gyrus (DG), cornu ammonis regions 1-3 (CA1-3), subiculum (SUB), entorhinal cortex (EnRC), Brodmann area 35 (BA35) or Perirhinal cortex (PRC), BA36 or ectorhinal cortex (EcRC) and parahippocampal cortex (PHC). For analyses, CA1-3 were combined into a single volume and CA4 and DG were combined into a single volume, henceforth referred to as CAs and DG, respectively. For extrahippocampal regions a normalized volume which was obtained by dividing the raw volumes by the number of slices in which the ROI appears was used as recommended in (Yushkevich et al., 2015).

#### Quality control

We performed thorough quality control measures on the data using both manual and automated approaches. We used Qoala-T (Klapwijk et al., 2019) and also visual inspection of the FreeSurfer output. We ran Qoala-T on all of our initial subjects. For those flagged by Qoala-T to be removed, we utilized FreeView to visualize and examine the FreeSurfer segmented brain images. We excluded cases that failed both Qoala-T and manual inspection, which resulted to the exclusion of 9 cases. It should be noted that T1w images of these cohorts are the pivotal aspect of the MRI, so they have already passed a number of quality control steps, and are often repeated on the day of the scan if the image quality is not sufficient for analysis (e.g. due to subject motion). For images that passed Qoala-T check, we randomly selected five subjects from each imaging site and manually inspected whether or not they should be excluded, based on the parameters set by FreeSurfer’s post-processing manual. No additional images were excluded at this stage. Furthermore, we performed an extensive manual quality control on the segmentations resulting from running ASHS by visualizing the segmented mask overlaid on the T2-TSE image (an example is presented in **Figure 1**). Images with failed miss-classification were excluded: 32 subjects were excluded due to failing ASHS quality control, due to low quality (n=8), incorrect T2-TSE field of view (n=4), or failed segmentation (e.g. outside hippocampus) (n=20), leading to the exclusion of total of 41 subjects. It should be noted that 63 subjects had two data time points, which were carefully checked using the quality assurance procedure described above and the time point with higher image quality was included in the study.

**Figure 1.**
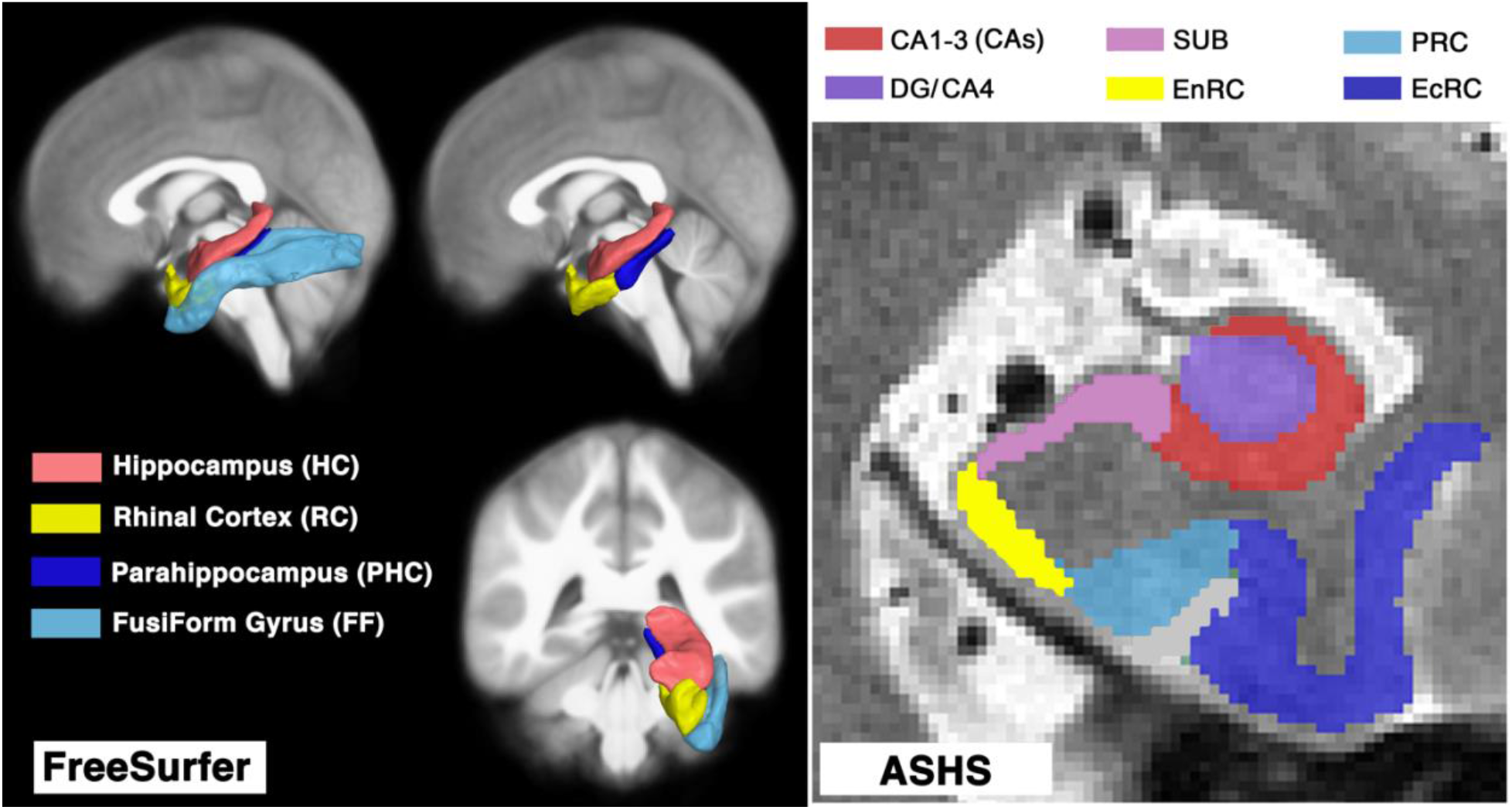
Schematic view of the medial temporal lobe regions of interest from FreeSurfer (left) and ASHS (right) software. The hippocampal formation, rhinal cortex, parahippocampal cortex and fusiform gyrus were segmented using isotropic T1-weighted (T1w) magnetic resonance imaging (MRI) with FreeSurfer’s atlas-based parcellation. Subfields of medial temporal lobe regions were then segmented using T1w and high resolution T2w MRIs with ASHS atlas-based parcellation. Given the parcellation uncertainty of smaller regions, dentate gyrus (DG) and cornu ammonis area 4 (CA4) were merged. CA1-3 were also merged into one region (referred to as CAs). Other parcellated regions are subiculum (SUB), entorhinal cortex (EnRC), Perirhinal cortex (PRC) and ectorhinal cortex (EcRC).

### Tau positron emission tomography

Tau positron emission tomography (PET) analysis was performed according to the UC Berkeley PET methodology for quantitative measurement (Baker et al., 2017; Landau et al., 2015, 2014; Schöll et al., 2016). Subjects were imaged by Flortaucipir (^18^ F-AV-1451). All PET images were co-registered on T1w MRI to align brain parcellation boundaries. Quantitative measurements were done based on standard uptake value ratio (SUVR). Tau PET includes a broad set of regional SUVRs, which includes cortical and subcortical regions and eroded hemispheric WM, which enables directed association analyses between regional volumetric values and corresponding Tau uptake.

### Cognitive scores in ADNI-3

#### Clinical dementia rating (CDR)

Six categories of cognitive functioning (memory, orientation, judgment and problem solving, community affairs, home and hobbies, and personal care) were assessed, in which the CDR describes five degrees of impairment (Hughes et al., 1982; Morris, 1993). Participant CDR scores were obtained through a standardized interview and assessment with the participant and a knowledgeable informant. Where a full CDR interview is not possible, the abbreviated CDR can be utilized.

#### Mini-Mental State Examination (MMSE)

The MMSE scale evaluates orientation, memory, attention, concentration, naming, repetition, comprehension, and ability to create a sentence and to copy two overlapping pentagons (Folstein et al., 1983). Participant MMSE score was obtained through standardized interview and assessment with the participant and a knowledgeable informant Participant categorization is detailed in the ADNI-3 protocol document. In brief:

#### Cognitively normal (CN)

Participants were considered CN if they had normal memory function, which was assessed using education-adjusted cutoffs on the Logical Memory II subscale from the Wechsler Memory Scale (Wechsler, 1987). Participants must also have an MMSE score between 24 and 30 and a CDR of 0 (memory box score must be 0). CN participants should be considered cognitively normal based on an absence of significant impairment in cognitive functions or daily living activities.

#### Mild cognitive impairment (MCI)

Participants were classified as MCI if they expressed subjective memory concern, with a score below the education-adjusted cutoffs on the Logical Memory II subscale from the Wechsler Memory Scale. The same MMSE score range as CN was used for the MCI group. Participants must have a global CDR of 0.5 and the memory box score must be at least 0.5. General cognition and functional performance of the MCI participants must be sufficiently preserved such that clinical diagnosis of Alzheimer’s disease cannot be made.

#### Alzheimer’s disease (AD)

Participants categorized as AD were excluded from this study, to ensure the focus remains on the early cognitive decline. In brief, participants were considered AD if they scored below the education-adjusted cutoff of Wechsler Memory Scale, had an MMSE score between 20 to 24 and global CDR>= 0.5 (Weiner et al., 2017).

### Cognitive scores in USC ADRC

Clinical Dementia Rating (CDR) assessments followed the standardized UDS procedures. Participants underwent clinical interview, including health history, and a physical exam. Knowledgeable informants were also interviewed. Participant CDR score was obtained through standardized interview and assessment with the participant and a knowledgeable informant.

### Statistical analysis

Tables 1 and 2 summarize study participant demographics and clinical information. When the normality hypothesis was rejected using D’ Agostino and Pearson’s test (Pearson et al., 1977), median and IQR were reported. Otherwise, mean and standard deviation were reported. Participant demographics and clinical characteristics were compared between CN and MCI groups using chi-square tests for categorical variables and one-way ANOVA for continuous variables. ADNI-3 and USC ADRC participants that were clinically diagnosed with Alzheimer’s disease were excluded from the analysis to keep the focus on early cognitive decline.

**Table 1.** Summary information of the participants of the discovery dataset (ADNI-3).

| Measures | Cognitively Normal | Mild Cognitive Impairment |
| --- | --- | --- |
| *n* | 429 | 175 |
| Female: *n* (%) | 59 | 42 |
| Age (yrs) | 73.19 ± 7.39 | 75.70 ± 8.12 |
| Education (yrs) | 16.93 ± 2.26 | 15.90 ± 2.80 |
| Brain volume (mm<sup>3</sup>) | 1,063,623 ± 106,788 | 1,052,296 ± 107,308 |
| CDR sum-of-boxes* | 0.0 ± 0.0 | 1.0 ± 1.5 |
| MMSE* | 29.0 ± 1.0 | 29.0 ± 3.0 |
| Ab PET (SUVR, AV45)<sup>+</sup> | 1.11 ± 0.17 | 1.16 ± 0.28 |
| HC Tau PET (SUVR, AV1451)<sup>++</sup> | 1.19 ± 0.15 | 1.27 ± 0.20 |
| RC Tau PET (SUVR, AV1451)<sup>+++</sup> | 1.10 ± 0.13 | 1.23 ± 0.25 |

Values are mean ± SD or frequency (percent).
\* values are median ± IQR (normality hypothesis was rejected with D'Agostino and Pearson's test)
+ n=235 subjects had AV45 Aβ PET data: n(CN)=177, n(MCI)=58
++ n=322 subjects had AV1451 tau PET data: n(CN)=244, n(MCI)=78
+++ n=378 subjects had data about history of hypertension: n(CN)=269, n(MCI)=86
CDR: clinical dementia rating, MMSE: mini-mental state examination, PET: positron emission tomography, SUVR: standard uptake value ratio, HC: hippocampus, RC: rhinal cortex

**Table 2.** Summary information of the participants of the replication dataset (USC ADRC)

| Measures | Cognitively Normal | Questionable or Mild Cognitive Impairment |
| --- | --- | --- |
| *n* | 121 | 54 |
| Female: *n* (%) | 81 (67%) | 25 (46%) |
| Age (yrs) | 68.94 ± 8.41 | 73.38 ± 7.73 |
| Brain volume (mm<sup>3</sup>) | 1,055,607 ± 110,713 | 1,048,289 ± 115,324 |
| Values are mean ± SD or frequency (percent). |  |  |

[here is a template:] We used linear regression when investigating the age-dependence of regional morphological atrophy using an ordinary least square fitting routine, implemented with the statsmodels.OLS module in Python 3.5.3 (StatsModels version 0.8.0 – other Python packages that were used are Pandas version 0.20.3 and NumPy version 1.13.1). Multiple regressions were fitted to regional volumes, one region at a time. For every instance, sex, age and years of education were included as covariates. The Benjamini–Hochberg procedure with a false discovery rate of 0.05 was used to correct for multiple comparisons. For each CN and MCI group, a morphometric correlation network was developed by calculating the cross-correlation similarity matrix in normal aging and early cognitive decline. Then, the absolute difference was calculated and shown in **Figure 7**.

## Results

Volume of the MTL regions decreased with age, with a fastest rate observed in HC (**Figure 2.a**). When overall age-related brain shrinkage was taken into account (measuring relative volume change of each MTL region), RC appeared to be intact in normal aging (**Figure 2.b**), while HC, FF and PHC showed a faster decline than overall brain aging (**Figure 2.c-d**). Relative volume of the RC was independent of age (*p*=0.97), while relative volume of the HC was significantly associated with age (*p*=1e-18). Relative volumes of FF and PHC were also associated with age, but in a smaller degree compared to HC (**Figure 2.b**).

**Figure 2.**
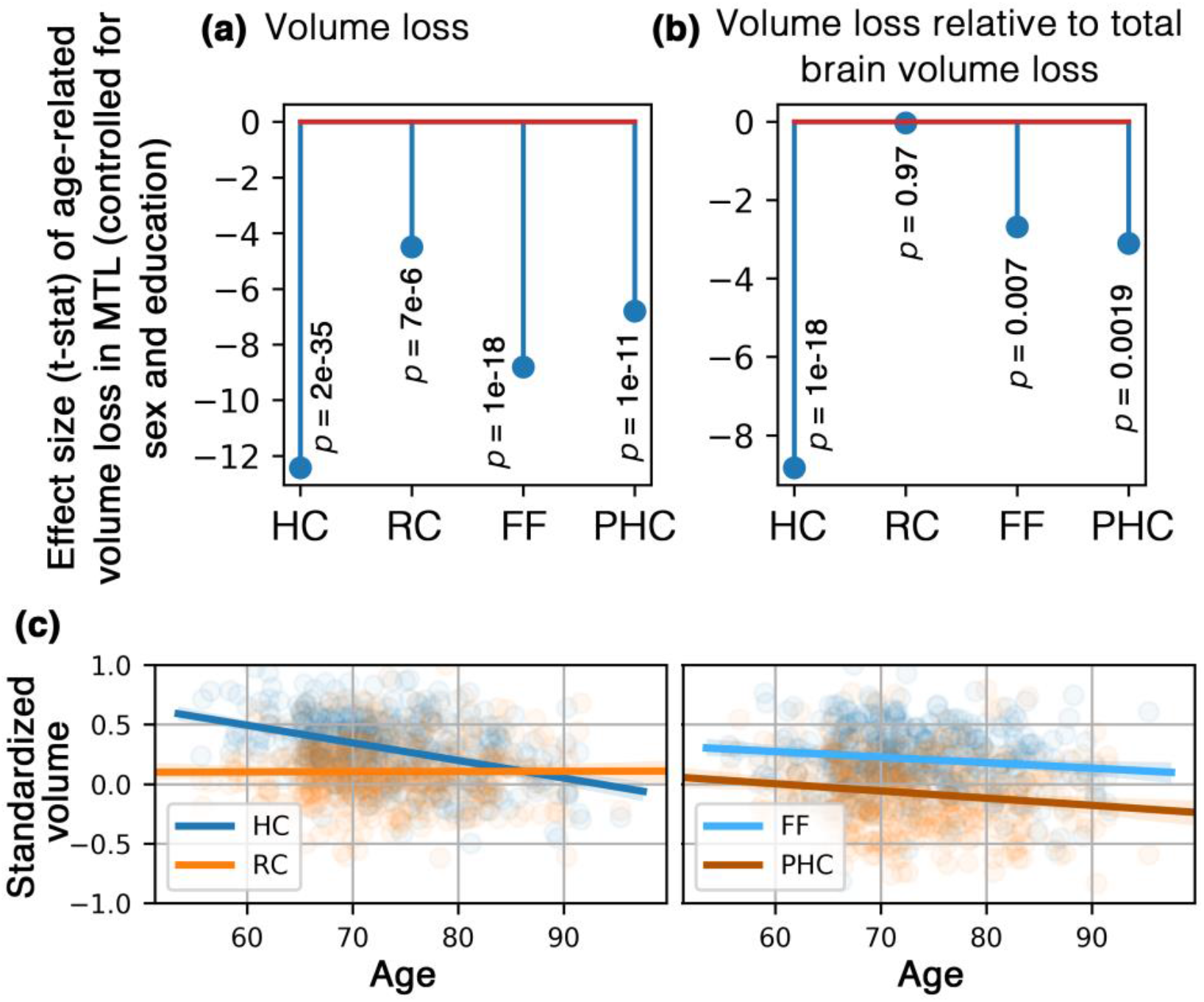
Age-dependent morphometry of medial temporal lobe (MTL) regions in cognitively normal aging participants (N=429). Age-related volume loss was observed in all MTL regions **(a)**. The rate of volume loss in the MTL regions exceeded that of total brain volume, with the exception of the rhinal cortex (RC) **(b)**. RC volume showed no age-dependency and remained intact in cognitively normal aging participants. Subplots showing the age-dependent volumetric changes across 4 regions of interest demonstrates the relative stability of the RC compared to the rest of the MTL regions **(c)**.

When ASHS output were used to explore subfields of MTL regions, we noted same age-dependency on the raw subregions volumes (**Figure 3.a**: volume decreased with age). However, when relative volume (volume divided by brain volume) were used, we noted that SUB, EnRC, PRC and EcRC were not significantly associated with age (**Figure 3.b-d**). In particular, EnRC relative volume appeared to be independent of age and remained intact (*p*=0.65). Relative volume of DG and CAs were age dependent and decreased with a faster rate than overall brain volume.

**Figure 3.**
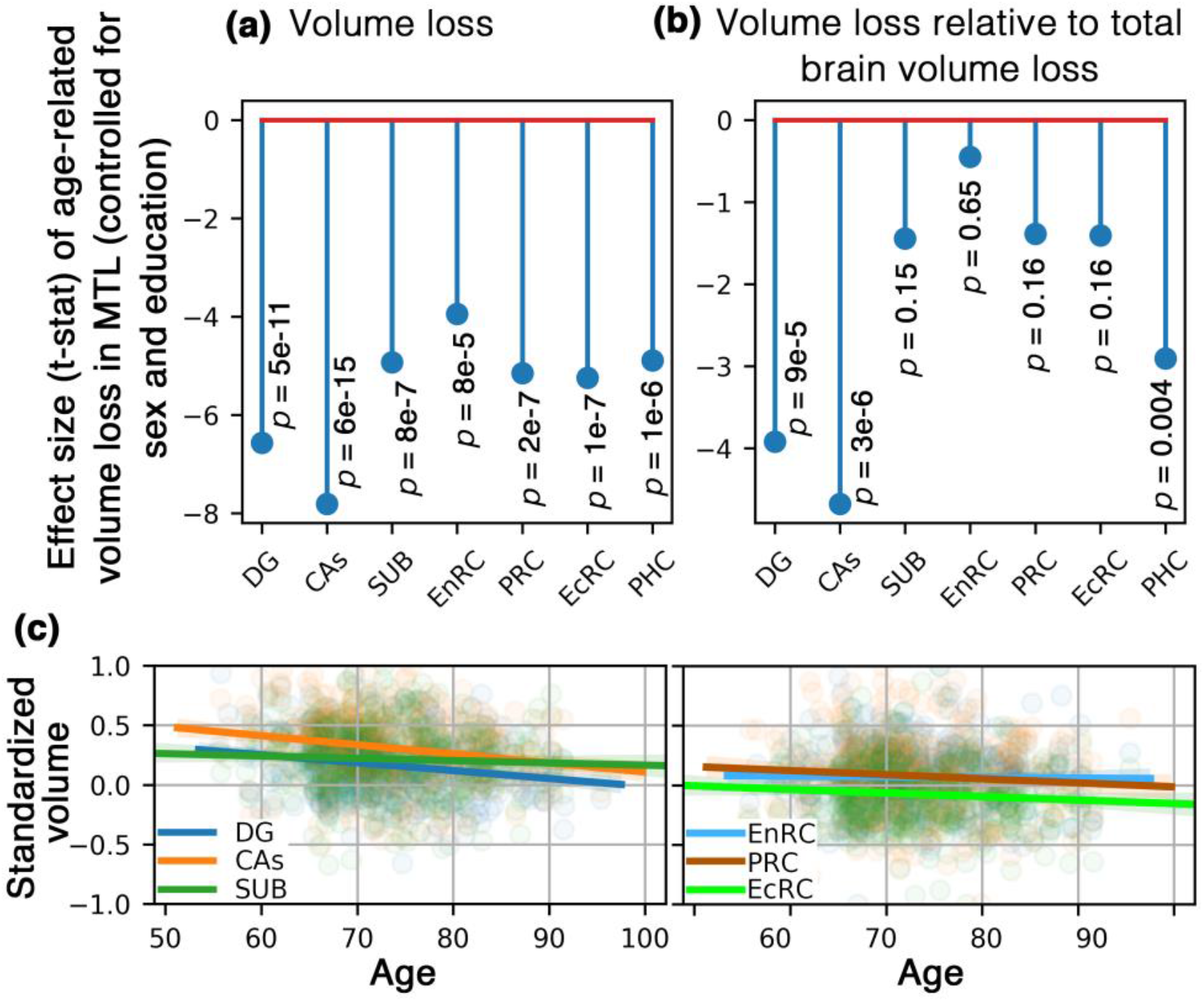
Medial temporal lobe subfield morphometry in normal aging participants (N=429). Age-related volume loss was observed in all MTL regions **(a)**. However, regions at the boundary of the hippocampal proper and rhinal cortex appeared to be preserved in cognitively normal aging subjects when total brain volume loss was considered **(b)**. Subplots showing age-dependent changes of MTL regions of interest standardized by total brain volume demonstrate entorhinal cortex (EnRC) is the least affected by the age-dependent atrophy, followed by the perirhinal (PRC) and ectorhinal (EcRC) cortices. Subiculum (SUB) of the hippocampal formation also remained relatively intact in comparison to DG and CAs.

When CN and MCI were compared, MCI participants had lower cortical volume across MTL regions, regardless of the age (**Figure 4**). We noted a similar age-dependency of cortical volumes in MCI and CN across all MTL regions, except for RC (MCI vs CN difference were highest between the older participants). Unlike CN participants, RC was not intact in MCI participants, which was the major age-dependent difference between CN and MCI. RC atrophy appeared to be a specific marker of the late-onset cognitive decline.

**Figure 4.**
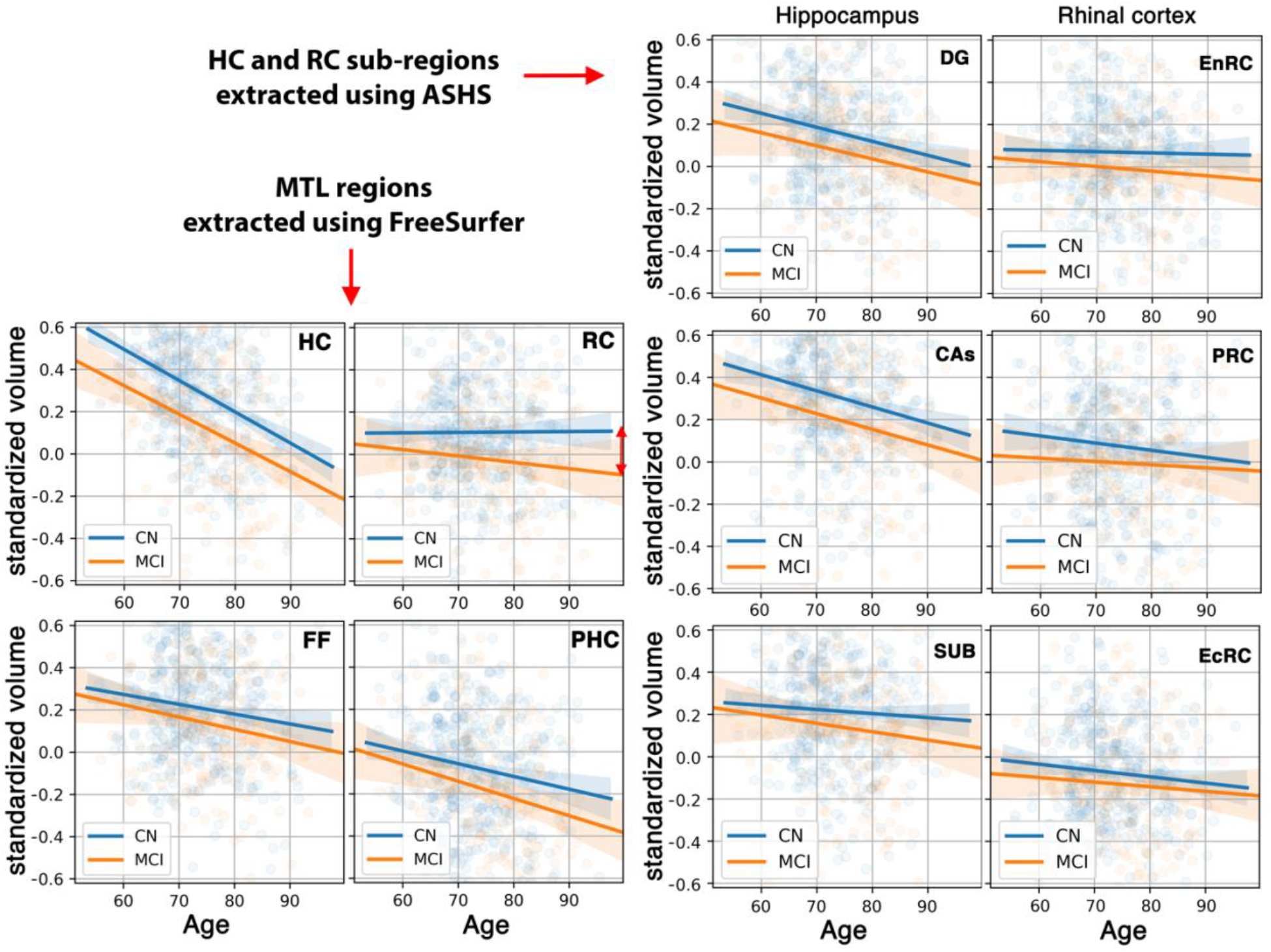
Age-dependent medial temporal lobe (MTL) volumetric differences, compared between cognitively normal (CN, N=429) and mild cognitively impaired (MCI, N=175) of ADNI3 participants. Values are standardized volume relative to total brain size. SUB and EnRC showed the least degree of age-dependent atrophy in CN. For these two regions, the largest difference between CN and MCI were observed in participants with late-onset cognitive decline.

When subregions of the MTL were investigated, SUB and EnRC showed similar results as RC, potentially explaining the localized focus of the observed RC age-dependency (**Figure 4**). In both SUB and EnRC, the MCI participants showed age-dependent atrophy, while the cortical volume of the CN group remained intact.

The replication dataset confirmed results of the discovery data (**Figure 5**), in which RC showed to be intact in CN participants (in the replication dataset, RC showed even a slower atrophy rate compare to overall brain ageing). Similarly, when subregions were looked at, transhippocampal regions (SUB, EnRC, PRC and EcRC) showed the biggest age-dependency difference between CN and MCI. Relative volumes of these regions were not associated with age in CN (particularly EnRC). The rate of atrophy was highest in EnRC in older adults with mild cognitive decline, suggesting that EnRC atrophy could be a specific biomarker of late-onset cognitive decline.

**Figure 5.**
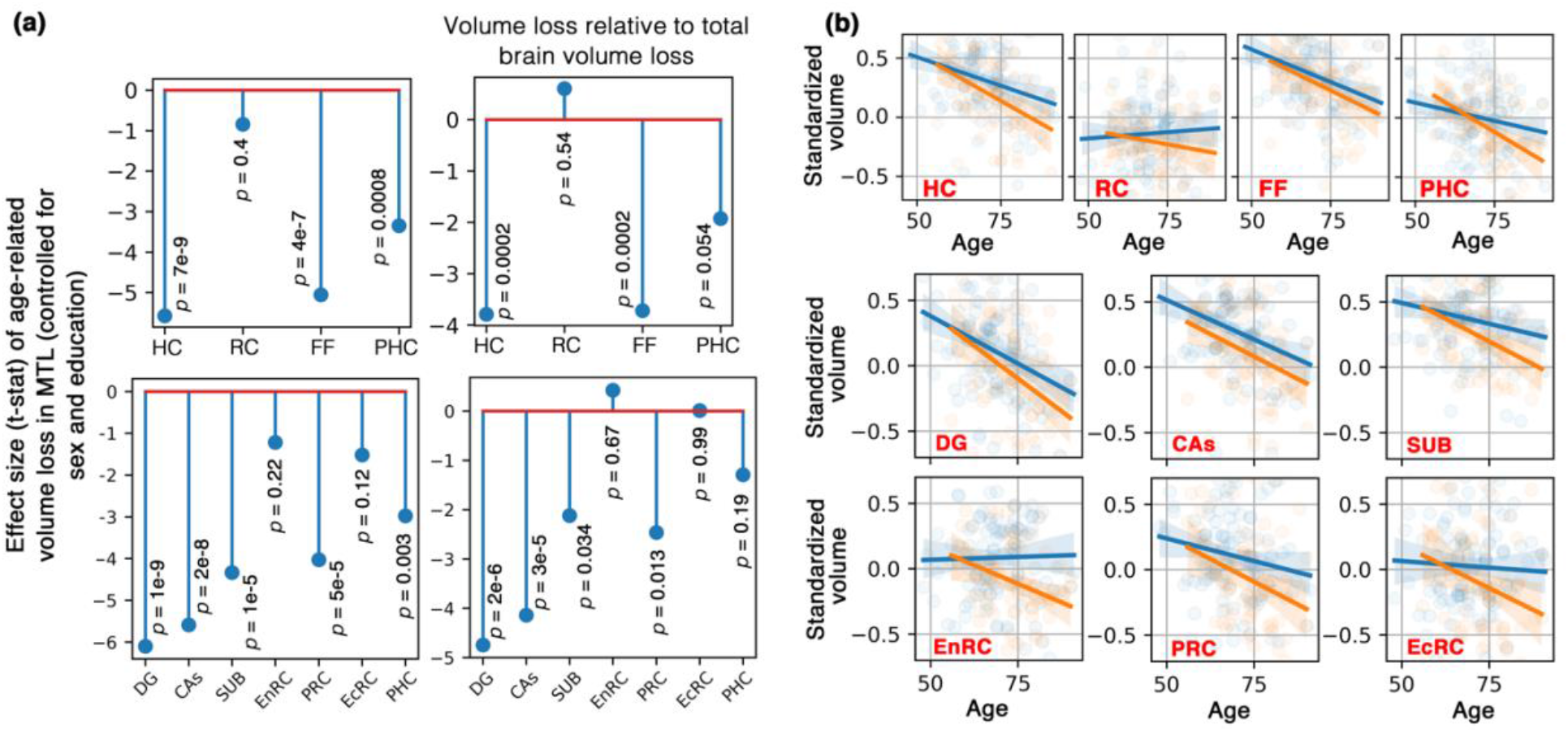
Replication study on the USC ADRC cohort (N=175). Age-dependence across regions of medial temporal lobe and their subfields across cognitively normal (CN) participants (N=121) are presented in **(a)**. Age-dependent volume difference of medial temporal lobe regions were compared between CN and mild cognitively impaired (MCI) participants (N=54) are presented in **(b)**; Blue: CN, Orange: MCI. In corroboration with previous findings, rhinal cortex (RC), and in particular entorhinal cortex (EnRC), appeared to be the key region in cognitively preserved aging.

Morphometric correlation network between DG-EnRC and CAs-EnRC were distinctly and most different in cognitively declined subjects compared to normal aging profile (**Figure 6**). A high value in the morphometric correlation network means that the pattern of atrophy in the network is distinct from normal brain aging profile. Corroborating previous analysis, transhippocampal regions, and in particular SUB, EnRC and PRC showed the most distinction from normal aging.

**Figure 6.**
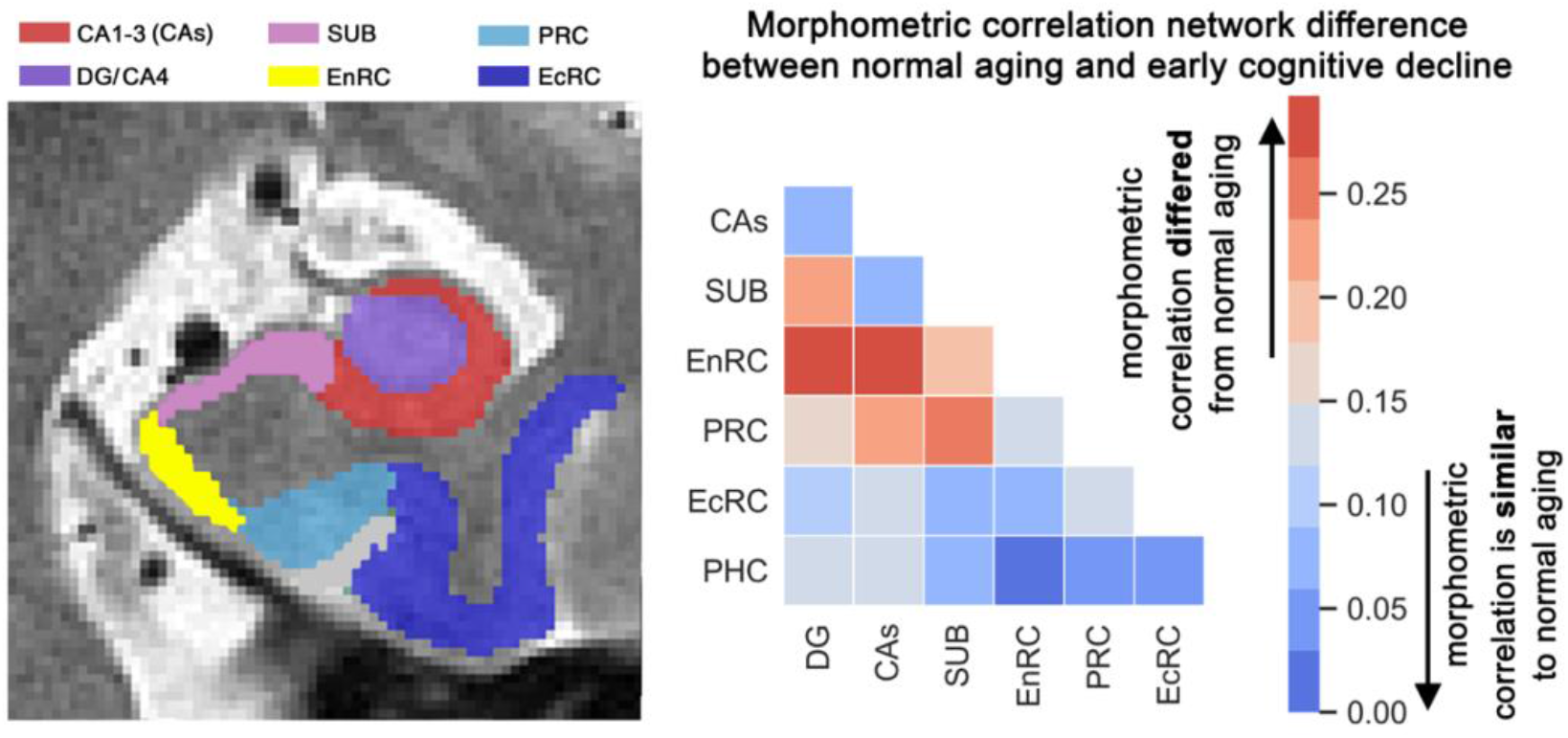
The morphometric correlation network. shows subiculum (SUB), entorhinal cortex (EnRC) and perirhinal cortex (PRC) were differentially affected by cognitive decline across age (N=604). The cognitive decline morphometric correlation matrix is the difference between the morphometric correlation matrix of normal aging participants (N=429) and that of cognitively impaired subjects (N=175).

**Figure 7.**
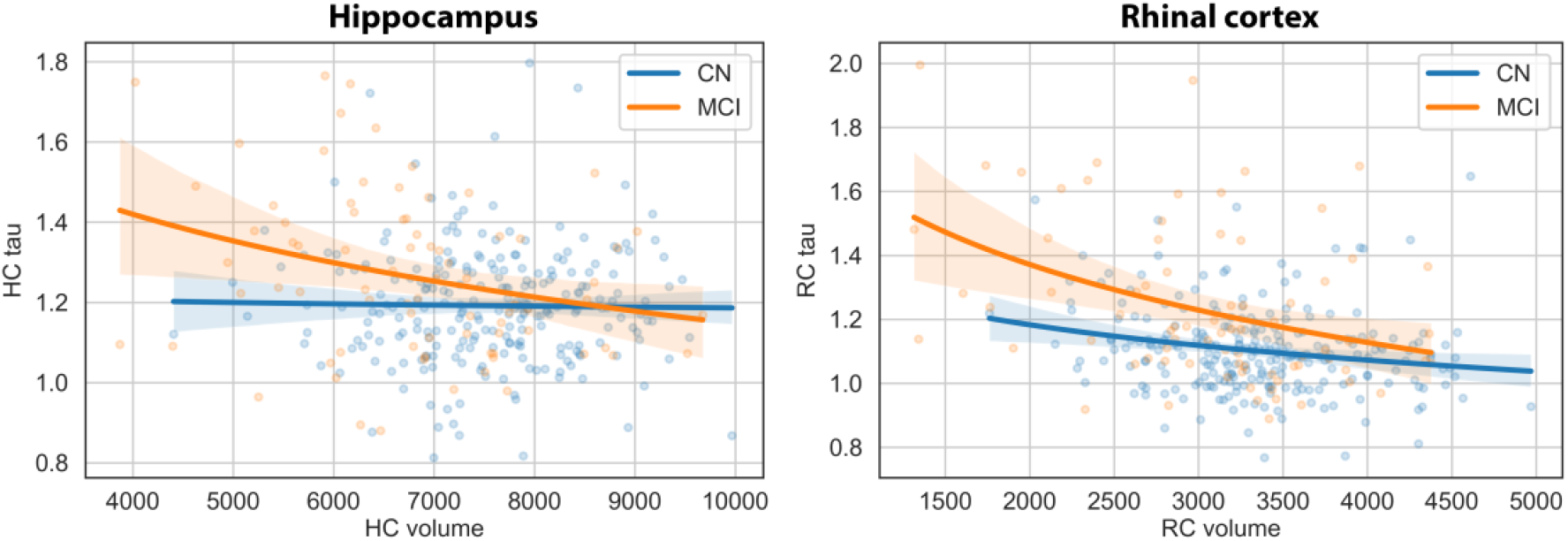
Correlation between regional volumes and tau uptake (N=322). Rhinal cortex (RC) and hippocampal (HC) atrophy were correlated with corresponding tau standardized uptake value ratios (SUVR), particularly in mild cognitively impaired (MCI) participants (N=78). Within cognitively normal (CN) participants (N=244), we observed a weak correlation between RC regional volume and tau uptake, which may reflect CN participants at risk of cognitive decline. This was not observed in HC, corroborating the expected Braak stage atrophy pattern of AD with early stage pathology observed in RC. This also shows that the combination of RC tau uptake and volume could have stronger diagnosis value in comparison to HC, which is commonly used for AD diagnosis.

A stronger correlation between regional volume and tau uptake was observed in RC compared with HC. Both regions showed a negative correlation between cortical volume and tau SUVR. Tau uptake and regional volume of MCI participants were significantly correlated in both HC (*b*(78) = −.00008, *t* = −3.9, *p* = .0002) and RC (*b*(78) = −.00012, *t* = −3.5, *p* = 0.0007). Within CN participants no correlation between HC volume and tau uptake was observed (*b*(244) = −.000015, *t* = −1.7, *p* = .08). However, we observed a significant correlation between Tau uptake and RC volume in CN participants (*b*(244) = −.000044, *t* = −3.0, *p* = .003), which may present CN participants with higher chance of converting to MCI. These results confirm that RC is a more specific target compared to HC. We generated similar figure including AD participants for sanity check, which corroborated the expected pattern (higher Tau uptake and lower regional volume – **Supplementary Figure 1**).

## Discussion

In the current study, we used T1-weighted and high-resolution T2-weighted images to assess age-related volumetric changes to medial temporal lobe regions, including the hippocampal formation and rhinal cortex, in patients with MCI and cognitively normal controls in order to differentiate between normative and pathological aging trajectories. Furthermore, findings were replicated in a second dataset and validated using tau PET measures as a pathological biomarker of neurofibrillary tangle deposition. Our results show age was significantly associated with regional atrophy in the hippocampus, but not the rhinal cortex. Additionally, variability in regional medial temporal lobe volume was associated with tau uptake in the rhinal cortex, but not the hippocampus. Because aging and AD pathology exert differential influences on MTL structure, these results may assist in the successful prediction of subsequent neurodegeneration and cognitive decline.

The influence of aging and neurodegeneration on hippocampal structure has been widely explored and several studies have demonstrated that hippocampal atrophy is a hallmark feature of both (Apostolova et al., 2012; Driscoll et al., 2009; Fjell et al., 2009; Jack et al., 1998; Pfefferbaum et al., 2013; Raz et al., 2010; Wang et al., 2006), however changes to the rhinal cortex have been less widely explored. In the present study, hippocampal structure undergoes significant reductions over time in both patients with MCI and cognitively normal controls. Specifically, atrophy was localized to the CAs and DG subfields, while the subiculum (SUB) was spared. In contrast, rhinal cortex structures were relatively preserved with age in cognitively normal older adults. Together, these results suggest that the rhinal cortex may provide more discriminative power for the early detection of MCI, as age-related hippocampal atrophy in cognitively normal subjects may obscure pathological changes in individuals converting to the preclinical stages of AD.

Previous studies have shown differential age-dependent vulnerabilities to subfield volume in cognitively normal older adults, with reductions observed in CA1 and DG subfield volumes with relative sparing of the SUB with age in cognitively normal older adults (Adler et al., 2018; Daugherty et al., 2015; Mueller and Weiner, 2009; Wisse et al., 2014). Studies in patients with MCI also mirror these changes, with significant changes observed in the CA1 (Chételat, 2018; de Flores et al., 2015b). These results are also in line with previous behavioral studies showing episodic memory decline with advancing age in the general population (Nyberg et al., 2012).

Diverse processes appear to be affected in hippocampal subfields during normative aging, and animal studies suggest volumetric reductions may be attributed to loss of pyramidal cells and granular cells in CA1 and DG, respectively (Driscoll, 2003), reduced dendritic complexity (Davies et al., 2003; Geinisman et al., 2004), axonal degeneration (Ypsilanti et al., 2008) and loss of glial processes (Hayakawa et al., 2007).

The relative age-invariance of the SUB in the current analysis was not observed in the replication study for either cognitively normal adults or patients experiencing cognitive decline, where age was significantly and negatively associated with SUB volume in normal aging adults and patients with MCI, thus underscoring the potential heterogeneity of aging effects on this hippocampal region. This could in part be due to the relatively smaller sample size of the replication study (n=175) compared to the discovery dataset (n=604). Lifespan studies have shown the SUB volume decreases steadily over time, though at slower rates compared to the rest of the hippocampal formation (de Flores et al., 2015b; La Joie et al., 2010; Malykhin et al., 2017; Ziegler et al., 2011). Furthermore, studies in patients with MCI also show significant changes to the SUB and surrounding areas (Carlesimo et al., 2015; Chételat, 2018; de Flores et al., 2015b; DeVivo et al., 2019). The discrepancies between our results and those shown in other studies may be attributed to varying segmentation strategies. It has previously been shown that the presubiculum is preserved in normal aging (Zheng et al., 2018), but undergoes initial atrophy during cognitive decline (Carlesimo et al., 2015). The segmentation protocol in the present study does not consider this demarcation; therefore, effects in the neighboring presubiculum may influence our results. Furthermore, other studies have shown that the molecular layer, the hypointense band containing neuropil between the DG and SUB, undergoes significant age-related changes in cognitively normal controls and patients with MCI (G A Kerchner et al., 2013) and the inclusion of this region will likely alter volumetric estimates of the subiculum. Our recent study using gene expression patterns suggest the SUB may be located more proximally along the hippocampal transverse axis than the ASHS segmentation indicates (Bienkowski et al., 2019).

In contrast to the age-related changes observed in the hippocampal formation, previous neuroimaging studies show that the effect of normal aging on the rhinal cortex is negligible (Juottonen et al., 1998; Raz et al., 2005, 2004; Su et al., 2018; Wang et al., 2019). Furthermore, microscopy studies in the entorhinal cortex demonstrate preservation of principal neurons with age in non-human primates (Gazzaley et al., 1997; Keuker, 2003; Merrill et al., 2000) and stability of membrane lipid composition in cognitively normal older adults (Hancock et al., 2017). The influence of age on rhinal cortex volume in patients with MCI, however, yielded mixed results where the replication study observed significant age-related atrophy in the entorhinal, perirhinal and ectorhinal cortices, which is supported by previous studies that have found significant differences in volume between patients with MCI and controls within the entorhinal cortex (Su et al., 2018; Trivedi et al., 2011) and perirhinal cortex (Wolk et al., 2017). The relative heterogeneity of aging trajectories in patients with cognitive decline compared to cognitively normal controls may be attributed to differences in time since disease onset, progression rate heterogeneity and variable manifestations of cognitive decline, as some patients with MCI will convert to AD while others will remain stable (Clem et al., 2017).

The extent of asymptomatic neurodegenerative pathology in the medial temporal lobe of cognitively normal and mildly impaired aging adults can be assessed *in vivo* with PET imaging (Cho et al., 2016; Maass et al., 2017; Schöll, 2015). Tau is a protein produced throughout the central nervous system that plays an important role in axon microtubule assembly and stabilization (Ballatore et al., 2007). Equilibrium shifts can lead to conformational changes in tau, resulting in a soluble hyperphosphorylated state that alters protein function, aggregates into neurofibrillary tangles (Kovacech et al., 2010; von Bergen et al., 2005) and is a hallmark pathological features of AD (Braak et al., 2006; Braak and Braak, 1996, 1991). Neurofibrillary tangle density can be detected decades before the onset of clinical symptoms (Younes et al., 2019) and is strongly associated with the severity of cognitive impairment and neurodegenerative processes as the disease manifests (Arriagada et al., 1992; Duyckaerts et al., 1987; Nelson et al., 2012; van Rossum et al., 2012). Tau pathology is initially observed in medial temporal lobe structures, where the spatial distribution of regional neurofibrillary tangle densities generally mirrors structural atrophy patterns during AD progression (Braak et al., 2006; Braak and Braak, 1996, 1991). Therefore, medial temporal lobe hyperphosphorylated tau is considered a strong biomarker for AD-related pathology. In the present study, we replicate previous findings of significant correlations between medial temporal lobe atrophy and tau uptake in patients with MCI. However, we also found regional tau deposition was significantly associated with atrophy of the rhinal cortex, but not the hippocampal formation, in cognitively normal aging adults. This is in contrast to our findings that hippocampal volume, but not rhinal cortex volume, is reduced with advancing age. Because a proportion of control subjects may eventually progress to MCI, rhinal cortex volume heterogeneity may be explained by early pathological changes consistent with AD. This is supported by histological (Braak and Braak, 1991; Liu et al., 2012) and *in vivo* neuroimaging evidence (Das et al., 2019) that rhinal cortex regions are uniquely affected by asymptomatic tau pathology before the clinical onset of dementia, prior to spreading to the hippocampal formation and the cortex. Together, these results suggest that tau deposition in the rhinal cortex may be a better indicator of conversion to MCI than the hippocampus due to the increased separation in aging trajectories between cognitively normal older adults and patients with MCI.

Interpretation of age-related medial temporal lobe structural differences in elderly adults is limited by the cross-sectional nature of the study. In contrast, longitudinal studies provide the opportunity to directly assess the influence of age on brain structures within subjects over time (Das et al., 2012; Driscoll et al., 2009; Fjell et al., 2009; Marcus et al., 2010; Parker et al., 2019; Raz et al., 2005, 2004; Resnick et al., 2003; Thambisetty et al., 2010). Cross-sectional studies can result in inconsistent results due to large inter-subject variability and methodological artifacts, such as cohort differences (Schaie, 2009), sometimes resulting in divergent and contradictory findings compared to longitudinal studies (Pfefferbaum and Sullivan, 2015). Relatedly, the present study may include cognitively normal older adults in the presymptomatic stage of dementia, which can result in an overestimation of the observed age-related decline (Burgmans et al., 2009).

In the present study, we used T1-weighted contrasts coupled with T2-weighted images with high in-plane resolution in clinically feasible acquisition times to identify sub-millimeter hippocampal subfields using automated segmentation protocols (Iglesias et al., 2015; Wisse et al., 2016). Several automated and manual segmentation strategies have been developed with different criteria for hippocampal subfield and cortical demarcations (Augustinack et al., 2013; Fischl et al., 2009; Iglesias et al., 2015; Pipitone et al., 2014; Van Leemput et al., 2009; Wisse et al., 2016; Yushkevich et al., 2015, 2010). The choice of segmentation method can alter the number and volume of regions, and different subfield combinations may make comparisons across studies difficult to assess. Interpretation of the present results in the context of previous research is therefore limited.

However, attempts are currently underway by the Hippocampal Subfields Group (HSG; http://www.hippocampalsubfields.com) to develop a unified segmentation protocol for medial temporal lobe regions (Wisse et al., 2017).

While the present study offers the ability to localize differences to sub-regions in the medial temporal lobe, we did not explore differences between anterior and posterior medial temporal lobe regions. The hippocampus exhibits a functional gradient along the anteroposterior axis (Poppenk et al., 2013) and previous studies have identified regionally specific and sometimes diverging effects of aging and neurodegeneration on regions along the long axis of the hippocampus (Malykhin et al., 2017; Martin et al., 2010; Yang et al., 2013). Future studies should aim to examine morphological changes within individual medial temporal lobe sub-regions to provide increased spatial specificity.

In conclusion, the results from this study demonstrate that age-related changes in medial temporal lobe structures in normal aging and MCI are localized to the hippocampus. Evidence from the PET study, however, indicates that tau uptake is associated with volume in the rhinal cortex of cognitively normal older adults. Therefore, rhinal cortex volume may be a better indicator of conversion to MCI than hippocampal volume because structures that are preserved in aging but affected in neurodegeneration may provide enhanced sensitivity for detecting early pathological changes in aging adults. These findings indicate that high-resolution MRI (Barisano et al., 2018; Sepehrband et al., 2018b) and regional volumetric measures of the medial temporal lobe can help identify structural biomarkers for the early identification of individuals at risk for developing AD. Future studies are needed to determine how the aging trajectory of the rhinal cortex and hippocampus change within subjects as they transform from normal aging to early cognitive decline in order to identify the inflection point separating normal and pathological aging.

## Acknowledgement

This work was supported by NIH grants: 2P41EB015922-21, 1P01AG052350-01 and USC ADRC 5P50AG005142. The content is solely the responsibility of the authors and does not necessarily represent the official views of the NIH.

## ADNI

Data collection and sharing for this project was funded by the Alzheimer’s Disease Neuroimaging Initiative (ADNI) (National Institutes of Health Grant U01 AG024904) and DOD ADNI (Department of Defense award number W81XWH-12-2-0012). ADNI is funded by the National Institute on Aging, the National Institute of Biomedical Imaging and Bioengineering, and through generous contributions from the following: AbbVie, Alzheimer’s Association; Alzheimer’s Drug Discovery Foundation; Araclon Biotech; BioClinica, Inc.; Biogen; Bristol-Myers Squibb Company; CereSpir, Inc.; Cogstate; Eisai Inc.; Elan Pharmaceuticals, Inc.; Eli Lilly and Company; EuroImmun; F. Hoffmann-La Roche Ltd and its affiliated company Genentech, Inc.; Fujirebio; GE Healthcare; IXICO Ltd.; Janssen Alzheimer Immunotherapy Research & Development, LLC.; Johnson & Johnson Pharmaceutical Research & Development LLC.; Lumosity; Lundbeck; Merck & Co., Inc.; Meso Scale Diagnostics, LLC.; NeuroRx Research; Neurotrack Technologies; Novartis Pharmaceuticals Corporation; Pfizer Inc.; Piramal Imaging; Servier; Takeda Pharmaceutical Company; and Transition Therapeutics. The Canadian Institutes of Health Research is providing funds to support ADNI clinical sites in Canada. Private sector contributions are facilitated by the Foundation for the National Institutes of Health (www.fnih.org). The grantee organization is the Northern California Institute for Research and Education, and the study is coordinated by the Alzheimer’s Therapeutic Research Institute at the University of Southern California. ADNI data are disseminated by the Laboratory for Neuro Imaging at the University of Southern California.

---

**Supplementary Figure 1.**
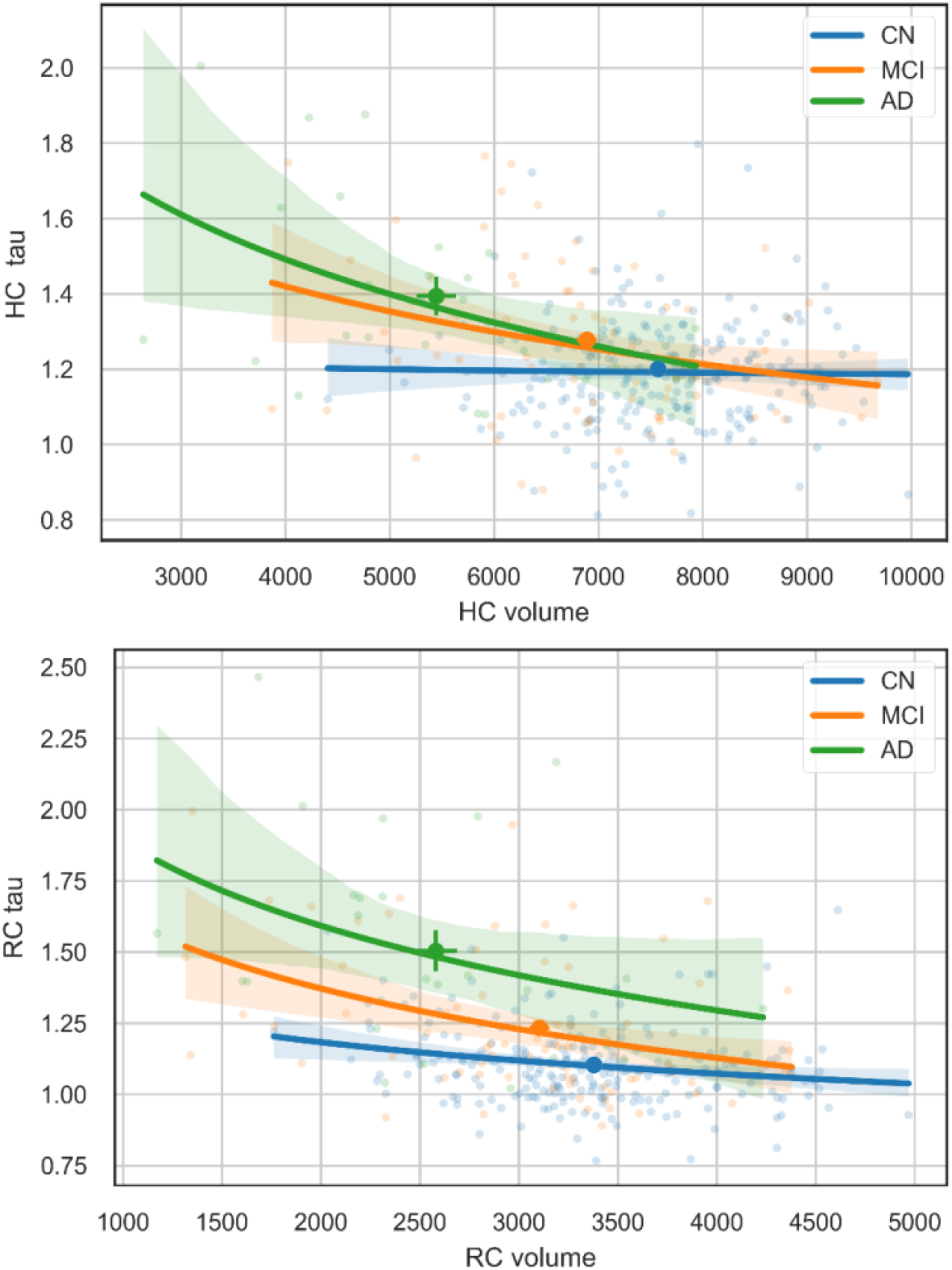
Correlation between regional volumes and tau uptake (N=348). Rhinal cortex (RC) and hippocampal (HC) atrophy were correlated with corresponding tau standardized uptake value ratios (SUVR), particularly in mild cognitively impaired (MCI) participants (N=78) and Alzheimer’s disease (AD) patients (N=26). Within cognitively normal (CN) participants (N=244), we observed a weak correlation between RC regional volume and tau uptake, which may reflect CN participants at risk of cognitive decline. This was not observed in HC, corroborating the expected Braak stage atrophy pattern of AD with early stage pathology observed in RC. This also shows that the combination of RC tau uptake and volume could have stronger diagnosis value in comparison to HC, which is commonly used for AD diagnosis.

